# Airborne point counts: a method for estimating songbird abundance with drones

**DOI:** 10.1101/2020.08.15.252429

**Authors:** McKenzie D. Somers, Darren B. Glass, Marisa A. Immordino, Precious S. Ozoh, Lauren B. Sherman, Andrew M. Wilson

## Abstract

Using drones to conduct airborne bioacoustic surveys is a potentially useful new way to estimate the abundance of vocal bird species. Here we show that by using two recording devices suspended from a quadcopter drone it is possible to estimate distances to birds with precision. In an experimental test, the mean error of our estimated distances to a broadcast song across 11 points between 0 and 100 m away was just 3.47 m. In field tests we compared 1-minute airborne counts with 5-minute terrestrial counts at 34 count locations. We found that the airborne counts yielded similar data to the terrestrial point counts for most of the 10 the songbirds included in our analysis, and that the effective detection radii were also similar. However, airborne counts significantly under-detected the Northern Cardinal (χ^2^_9_ = 22.8, post-hoc test *P* = 0.007), which we attribute to a behavioral response to the drone. Airborne counts work best for species that vocalize close to the ground and have high frequency-range songs. Under those circumstances, airborne bioacoustics could have several advantages over ground-based surveys, including increased precision, increased repeatability, and easier access in difficult terrain. Further, we show that it is possible to do rapid surveys using airborne techniques, which could lead to the development of much more efficient survey protocols than are possible using traditional survey techniques.

**Lay Summary:**

- We show that it is possible to estimate the distance of singing birds from a drone, which then allows bird counts to be converted to true abundance or population densities.
- Using drones to count birds allows researchers to survey areas that may be difficult or dangerous to access on foot.
- Airborne counts are potentially a highly efficient and highly repeatable way to estimate populations of vocal bird species.

## Introduction

Drones are now widely used in ecology to locate, count, or track organisms, and to map resources (Nowak et al. 2019). Most studies that have used drones to count birds have focused on larger species such as seabirds (Hodgson et al. 2018) and waterbirds (Afán et al. 2018, Pöysä et al. 2018) which are more likely to be detected in aerial imagery. The use of drones for bioacoustic surveys is less well-established, but initial studies have shown promising results for songbirds (Wilson et al. 2017) and bats (Kloepper and Kinniry 2018, Fu et al. 2018, August and Moore 2019). Drone-based bioacoustic studies have the potential to harness the advantages of bioacoustic techniques using automated recording units (ARUs) (Darras et al. 2018), together with increased mobility and ease of access in difficult terrain. Increased access could reduce habitat biases that are prevalent in bird survey data (Betts et al. 2007, Leitão et al. 2011). Other advantages of ARUs include obtaining a permanent record that can be analyzed/reanalyzed at a future date, and reducing observer bias (Campbell and Francis 2013, Shonfield and Bayne 2017). Further, while it is true that drones can cause disturbance to wildlife (Mulero-Pázmány et al. 2017), drones also have the potential to reduce disturbance caused by field biologists wandering through sensitive habitats (Christie et al. 2016, Borrelle and Fletcher 2017).

A previous study found that using inexpensive recording devices suspended from quadcopter drones to count songbirds produced counts that were broadly similar to those obtained by typical ground-based point count protocols (Wilson et al. 2017). However, the aim of many bird survey techniques is to estimate bird abundance, usually standardized to a given number of individuals per unit of area (Bibby et al. 2000), which then allows more direct comparison across species, locations, habitats, or time (Gregory et al. 2004). For airborne bioacoustic techniques to provide a useful alternative to ground-based survey methods (e.g. point counts, transects, territory mapping), the ability to estimate abundance within a given spatial area is highly desirable.

Here, we show that it is possible to estimate radial distances of bird vocalizations from a drone using two small inexpensive recorders, and a prosumer-grade quadcopter. We utilize the time difference of arrival technique (TDoA) (Mennill et al. 2006) to estimate distances to vocalizing birds on a horizontal plane either at ground-level, or at given heights above the ground. Estimation of distances to birds would allow the application of Distance Sampling techniques (Buckland et al. 2005) to estimate population densities. We show an experimental proof of concept and a field application, where we compare density estimates from airborne counts with those of traditional terrestrial point counts and territory/spot-mapping (Bibby et al. 2000).

## METHODS

### Equipment

We used a DJI Mavic Pro quadcopter programmed to fly missions autonomously using the Litchi app (VC Technology Ltd.) for iOS on an iPhone SE. We used aftermarket “Low-noise” DJI propellers, which reduce drone noise by ∼4 dB (Valle and Scarton 2019). Our recording devices were lightweight and inexpensive Zoom H1 Handy recorders, which weigh just 95 g, including battery, microSD card and, importantly, a windshield. It should be noted that this model of recorder was also chosen because of its cardioid pickup pattern, which reduces sound pickup from the rear of the microphones, i.e. in the direction of the drone when the microphone is directed at the ground. We attached the recorders to the drone using a system of fishing line, zip ties, and small carabiners.

### Measuring distances

To enable the use of time difference of arrival to estimate distances, the recorders had to be sufficiently far apart that time differences were measurable, and they had to be suspended below the drone to reduce excessive drone noise on the recordings. Previous tests showed that suspending recorders up to 15 m below the drone was manageable in the field, where great care is required to avoid entangling the fishing line on the drone or vegetation. Our system placed one recorder 7.5 m below the drone, and the second recorder 15 m below the drone. To calculate TDoA, the two recorders needed to be time-synchronized. We did this manually by placing the microphones of the two recorders ∼2 cm apart and playing a tree cricket (*Oecanthus* sp.) recording from an iPhone placed between them, thereby allowing the two recordings to be clipped to the same time point (to <1 mS accuracy) in Audacity (Audacity Team 2019). We then merged the two recordings in Audacity to make a single stereo audio track, which included time-differences in sound sources. TDoAs were measured manually from spectrograms (Hanning window with 512 samples and 89% overlap) in program Raven Pro 1.5 (Bioacoustics Research Program 2014).

With TDoAs measured we were then able to apply the Pythagorean Theorem to calculate the radial distance from the drone location to the sound source (*x*), in meters across the ground, using the following formula:

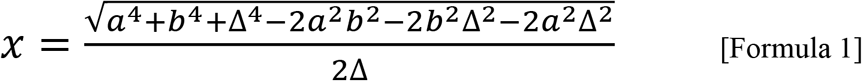

where *a* is the altitude of the bottom recorder (in meters), *b* is the altitude of the top recorder (meters) and Δ (Δ_cd_ in Figure 1) is the estimated difference in the Euclidean distance from the recorders to the sound source (*c-d*). Δ is calculated by multiplying the TDoA by the speed of sound at a given air temperature (*SOS_t_*). Hence, for any given TDoA, we could estimate the radial distance (*x*) between the point under the drone and a sound source.

**Figure 1.**
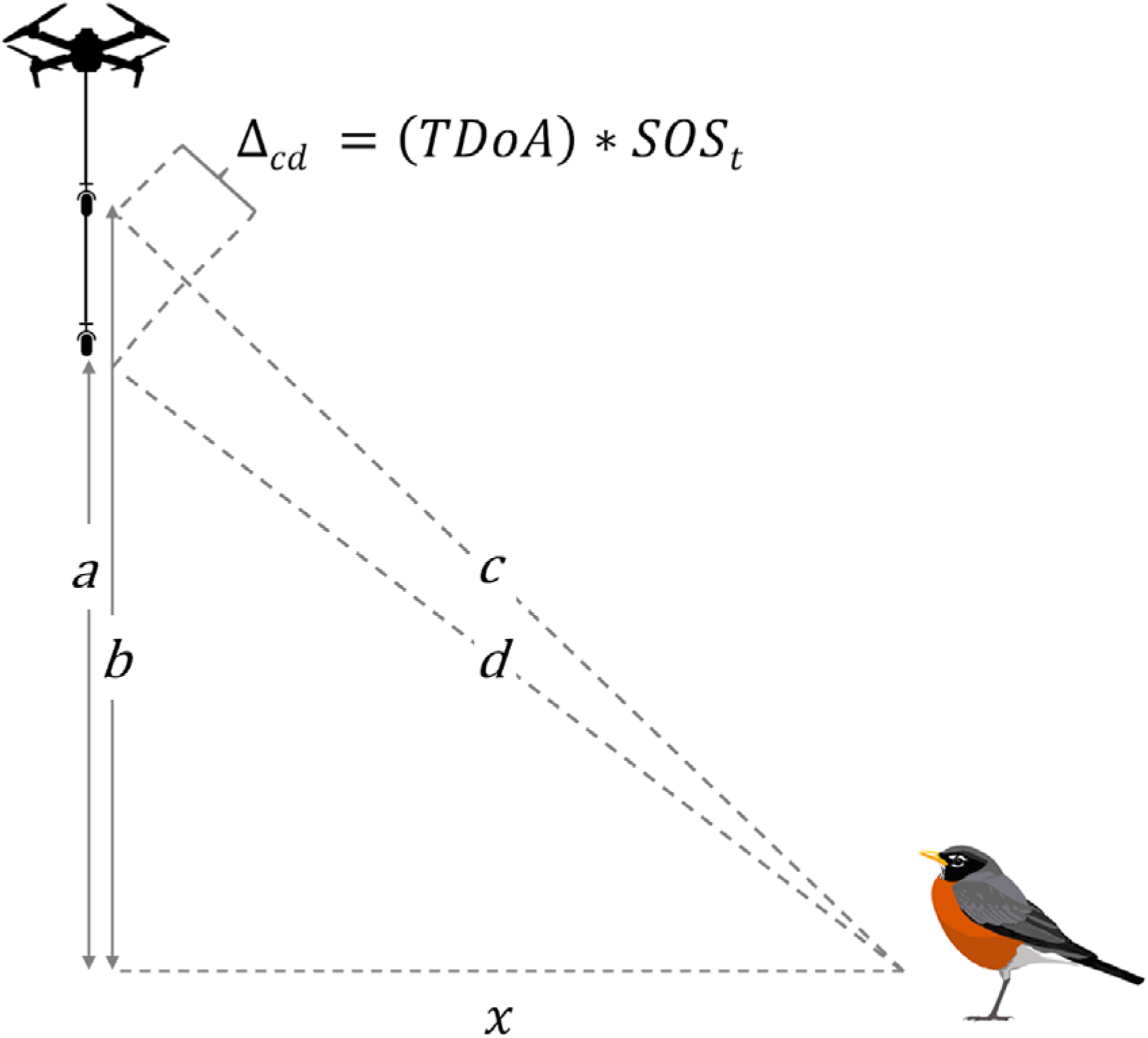
Application of the Pythagorean theorem to estimate radial distance from the location under a drone to a bird (*x*), from known heights of two recorders from the ground (*a* and *b*), and Time Difference of Arrival (*TDoA*) of a bird vocalization at recorders, based on speed of sound at a given air temperature (*SOS_t_*) Δ_cd_ is *c*-*d*, where *c* and *d* are unknown distances, to be estimated using Formula 1.

Because we were only able to measure time difference to whole mS (the measurement limit in Raven Pro), we must assume that the actual time difference was in a range of the measured TDoA ± 0.5 mS. Hence, we calculate the distance of the bird to be in a range between a lower (*x_l_*) and upper (*x_u_*) limits, and assuming that birds are randomly distributed within the band between those distances, a single estimate can be derived by estimating the median distance between the two:

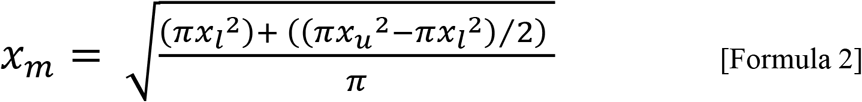

*x_m_* is therefore the estimated radial distance to the bird, which can used in distance sampling, or fixed radius distance abundance estimation from airborne point counts.

We used MS Excel to estimate radial distances to the sound source based on the above formulae. A spreadsheet that allows the user to estimate distance from measured TDoAs based on inputted air temperature (*t*), and recorder heights (*a*, *b*) is provided in Appendix 1. It is important to note that Formula 1 assumes that the sound source is on a horizontal plane at ground level. However, this assumption can be changed, for example, it can be assumed that a certain species typically sings from vegetation 4 m above ground-level, hence, heights *a* and *b* would be reduced by 4 m. Height of bird from the ground is therefore included in the spreadsheet as an additional parameter, that can be varied according to species and habitat. To test how sensitive the technique would be to uncertainty in estimating the height of birds from the ground we assumed an air temperature of 20°C and recorder heights at 40 and 47.5 m, and varied the height of the bird from the ground parameter from 0 to 10 m. We also used the spreadsheet to evaluate the effects of using different distances between the two recorders on TDoAs, which might be varied according to the study species and habitats, and research needs.

### Experimental test

To test the method we broadcast a bird song recording (American Robin *Turdus migratorius;* source: Macauley Library. 2014) at 90 dB at 1 m (measured using an Extech 407730 sound level meter) from paired Aomais Go speakers, placed on the ground. We then flew a mission where the drone hovered for 1-minute at altitudes of 55 m at 11 locations in 10 m increments along a transect, 0 m to 100 m across the ground from the broadcast speakers. The experiment was conducted on sports fields at Gettysburg College, PA, where background anthropogenic and biological noise was minimal. The temperature was 21°C and winds were light.

### Field test

With proof of concept established in our experiment, we sought to test the method in a field study, where we compared it to two traditional songbird survey techniques: point counts, and territory/spot mapping (Bibby et al. 2000). These survey methods are henceforth referred to as “mapping” and “terrestrial counts”, and surveys using the drone are referred to as “airborne counts”. The study was conducted in a section of State Game Lands 249, Adams County, Pennsylvania; 140 hectares of grassland and shrubby fields, with some small woodlots and wetlands (39.9374°N,-77.1774°W). Two tracks provide vehicular access to the site, but otherwise, the area has little human disturbance. Recreational drone flying is not permitted at the site. We focused our surveys on ten songbird species known to be present in sufficient numbers (>30 individuals): the Willow Flycatcher (*Empidonax traillii*), House Wren (*Troglodytes aedon*), American Robin (*Turdus migratorius*), Field Sparrow (*Spizella pusilla*), Song Sparrow (*Melospiza melodia*), Eastern Towhee (*Pipilo erythrophthalmus*), Yellow Warbler (*Setophaga petechia*), Common Yellowthroat (*Geothlypis trichas*), Northern Cardinal (*Cardinalis cardinalis*), and Indigo Bunting (*Passerina cyanea*).

For the airborne and terrestrial point counts we surveyed 34 pre-determined locations evenly spaced on 200-meter grid (data from a 35^th^ point were dropped due to excessive wind noise on drone recordings). Terrestrial and airborne counts were conducted on the same day, between 31^st^ May and 6^th^ June 2019, and between 6:00am and 9:00am. Weather conditions were suitable for both count methods, i.e. no rain, and wind less than a force 4 (15 mph) on the Beaufort scale (NOAA). Terrestrial counts were of 5–minute duration, with observations categorized by minute of first detection. Estimations of distances to birds seen or heard were aided by a laser range finder. Bird detections were categorized as singing, calling, or visual only. Terrestrial counts were all conducted by a single very experienced point count surveyor (A.M.W.).

For airborne counts, we hovered the drone at 55 m above ground-level for 1-minute. As in the experimental test, the two recorders were suspended 7.5 m and 15 m below the drone, hence, the recorders were 40 m and 47.5 m above ground-level. The drone was launched from at least 100 m away from the count locations. We conducted between two and five adjacent airborne counts in succession, which was readily done on a single battery. Based on personal observation and understanding of the habitat at our site, we estimated that most of the singing birds were in low vegetation. In our distance calculations we estimated that Willow Flycatcher, Field Sparrow, Song Sparrow, and Common Yellowthroat were 2 m above ground level, House Wren and Yellow Warbler 3 m, Eastern Towhee 4 m, American Robin and Northern Cardinal 5 m, and Indigo Bunting 6 m. In program Raven we labeled each bird on the airborne count recordings with a unique code so that we could track song output and possible movement for each individual, based on song-bout spacing, unique song patterns, apparent volume (from spectrograms) and calculated distances. We found very few instances of ambiguity when it came to identifying individual birds, based on these factors, even for the most abundant species.

To provide context for our point count data we conducted a territory mapping study of the entire site, visiting all areas on at least four occasions between mid-May and early July 2019. While the optimal number of visits for territory mapping is 10, a minimum of 4 will suffice (Gregory et al. 2004). In order to ensure the entire study area was surveyed as quickly as possible, teams of researchers surveyed different sections of the study area simultaneously. We estimated the number of territories using standard protocols (Bibby et al. 2000), with a lower estimate where territory clusters needed observations on two visits at least 10 days apart, and a higher estimate where single visit detections were also included as territories.

### Analytical methods

We estimated the density of singing birds from the terrestrial and airborne count data using the R package *Rdistance* (Miller et al. 2019). Half-normal, Hazard Rate and Negative Exponential detection functions were fitted and the best model was selected using AIC. Density estimates were compared with the estimated number of territories to see whether there was broad agreement in abundance estimates among the three bird survey techniques. Calculated Effective Detection Radii for terrestrial and airborne counts indicate whether the airborne counts are able to capture song detections over a similar area to a fieldworker on the ground. We tested to see whether there were differences in the species make-up of detections between airborne counts and 5-minute terrestrial counts using chi-square tests and post hoc tests of residuals with Bonferroni adjustment, using R package *chisq.posthoc.test* (Ebbert 2019)

## RESULTS

### Experimental results

Our experimental test showed that we were able to estimate the distance to the American Robin song broadcast with a high degree of accuracy; the mean absolute error was 3.47 m, which resulted in a mean overestimate of distances of 1.9 m (Figure 2).

**Figure 2.**
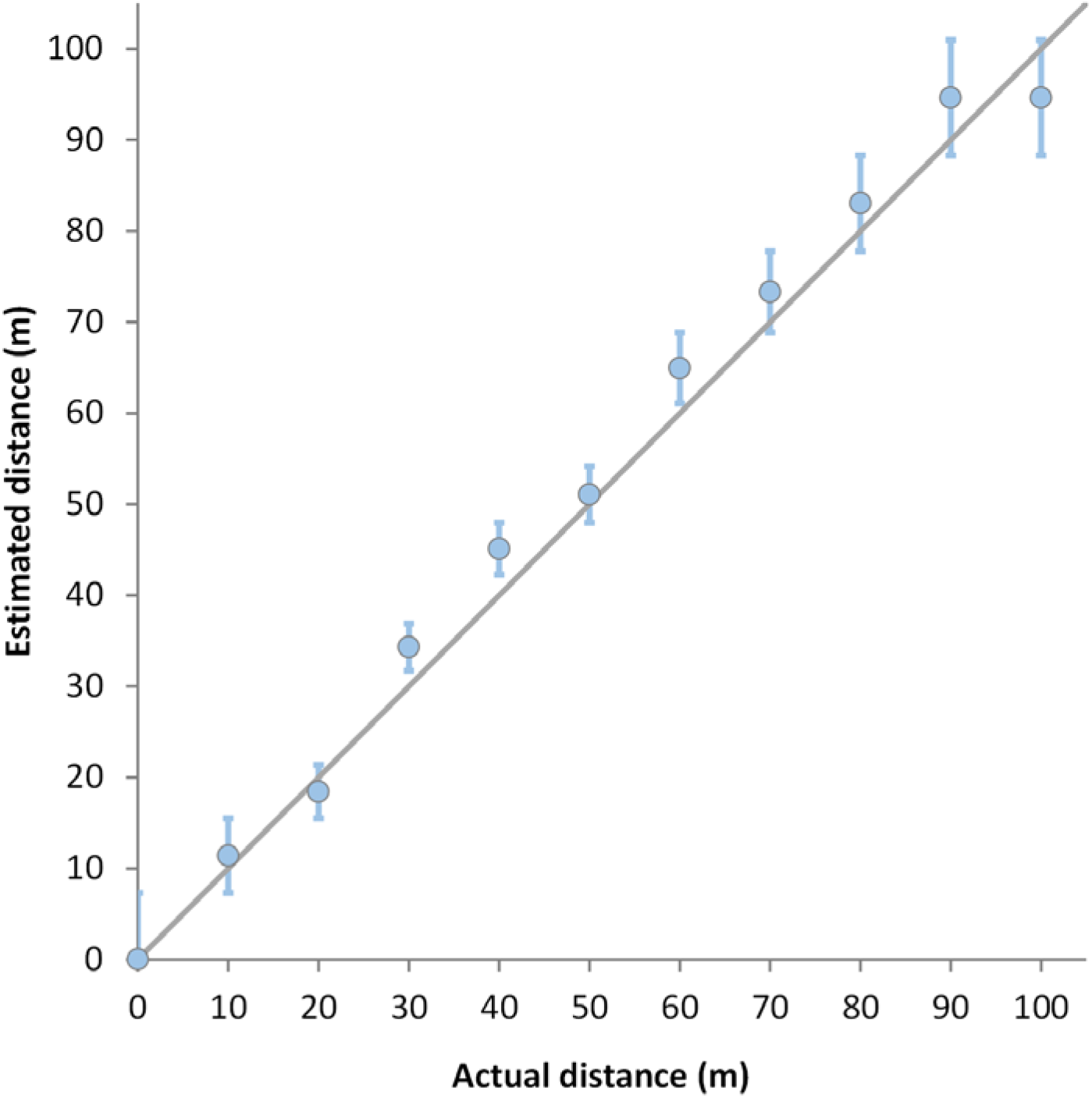
Actual versus estimated radial distances from a location under a drone flown at 55 m, to a broadcast American Robin song at ground level. Error are range bars.

We found that varying the distance between the two recorders affects the precision with which TDoAs can be measured. We note that the TDoA is maximized when the sound is directly below the drone, in which case the TDOA is given by *SOS*\*(*b*-*a*). In particular, if the recorders were 5 m apart, the maximum feasible TDoA at an air temperature of 20°C is 15 mS, compared with 22 at 7.5 m, and 29 at 10 m. More generally, one can directly show from the formula that as the distance between recorders is increased the formula for *x* becomes less sensitive to errors in the measured value of TDoA (range bars in Figure 3A).

**Figure 3.**
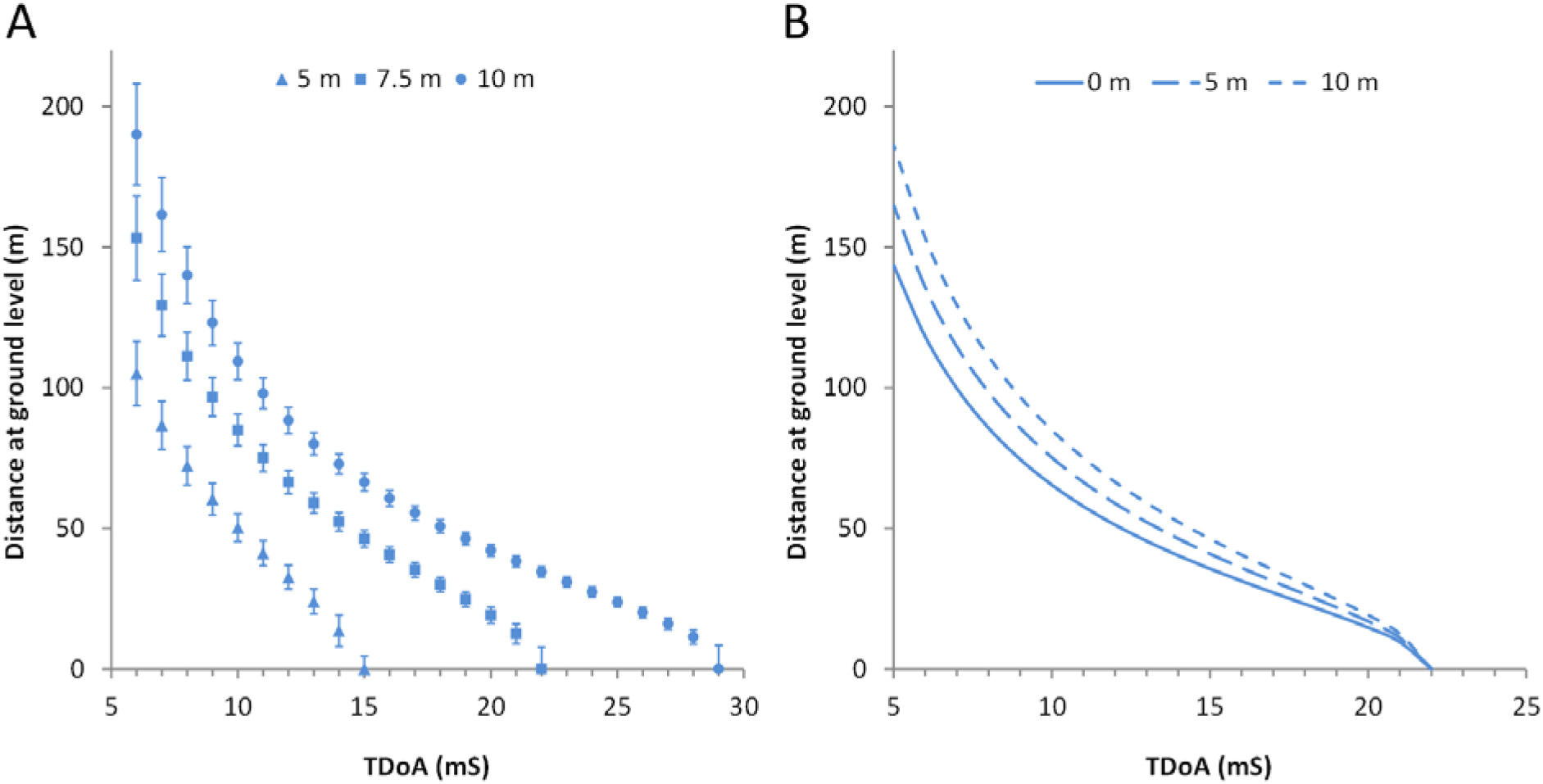
A: hypothetical radial distance estimates for measured TDoAs for three different distances between two recorders, assuming an air temperature of 20°C and drone altitude of 55 m. B: hypothetical estimated radial distances if it is assumed that a sound source is at three different heights off the ground.

The same computation shows that raising the height of the drone, while maintaining the distance between the two recorders constant, will make the formula less sensitive to error calculations. Uncertainty with respect to the height of birds off the ground will have rather little effect on distance estimation for birds close to the drone, but estimates would be increasingly uncertain at greater distance (Figure 3B). For example, if a TDoA of 10 mS is measured, the estimated distance to a bird on the ground is 85.3 m, compared with 75.6 m if it is assumed the bird is 5 m off the ground, and 65.8 m if the bird is 10 m off the ground.

### Field test

We detected 603 song bouts across our 10 target species on the 34 airborne counts; of which 369 song bouts were estimated to be by birds within a 100 m radius of the point location (Table 1). Detections beyond 100 m were discarded from further analysis due to the possibility of duplication between adjacent points. We found that song output was consistent throughout the one-minute airborne counts, with no indication of a curtailment of song activity in the drone’s presence (Figure 4A). We attributed the 369 song bouts to 120 different individuals (Table 1), of which 73% were detected within the first 20 seconds (Figure 4B).

**Table 1.**
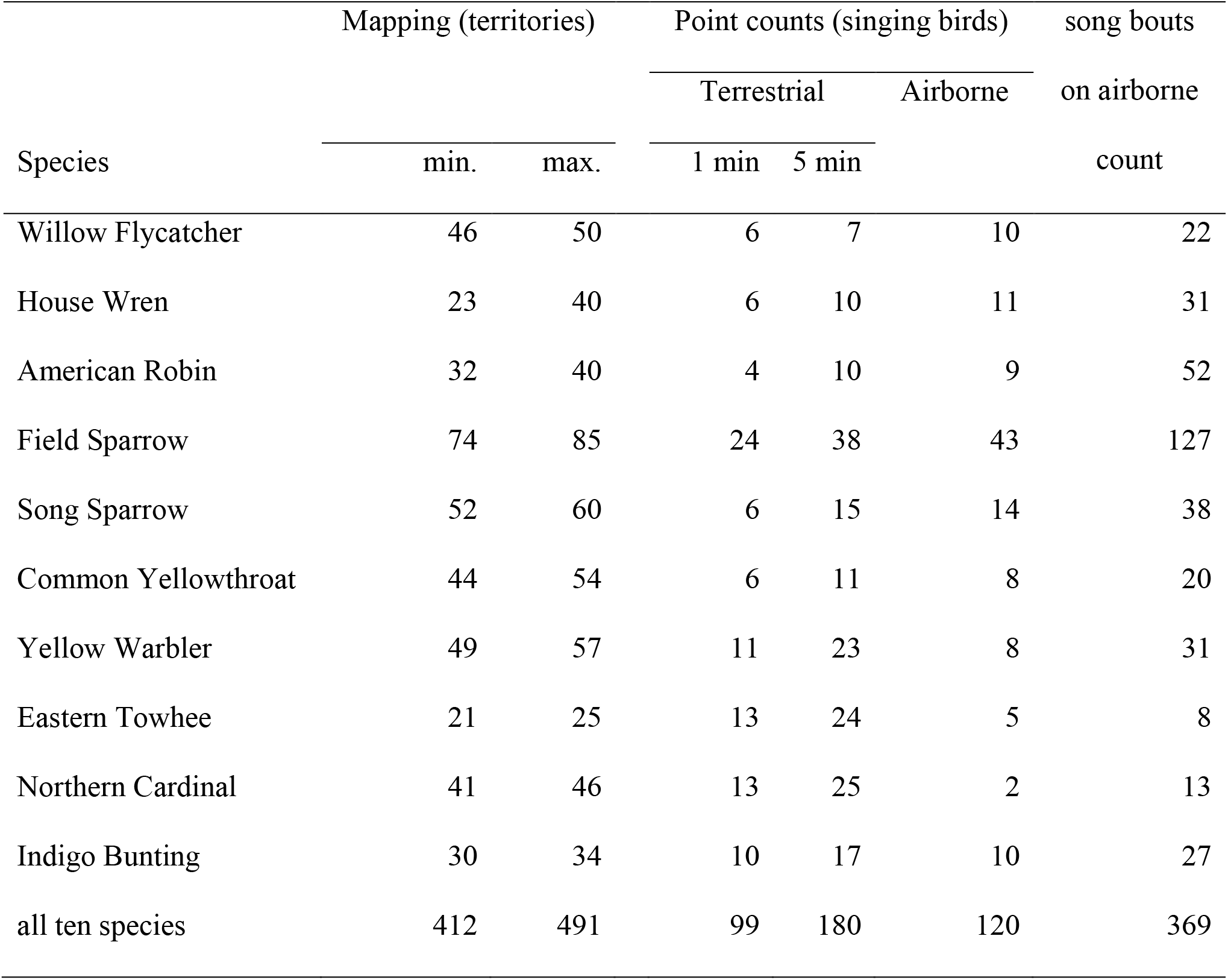
Estimated number of territories and point count detections (of singing birds) within 100 m of the point location, for terrestrial counts and airborne counts, 136 ha study area of State Game Lands 249, Pennsylvania, in 2019.

| Species | Mapping (territories) |  | Point counts (singing birds) |  |  | song bouts |
| --- | --- | --- | --- | --- | --- | --- |
|  | min. | max. | Terrestrial |  | Airborne | on airborne |
|  |  |  | 1 min | 5 min |  | count |
| Willow Flycatcher | 46 | 50 | 6 | 7 | 10 | 22 |
| House Wren | 23 | 40 | 6 | 10 | 11 | 31 |
| American Robin | 32 | 40 | 4 | 10 | 9 | 52 |
| Field Sparrow | 74 | 85 | 24 | 38 | 43 | 127 |
| Song Sparrow | 52 | 60 | 6 | 15 | 14 | 38 |
| Common Yellowthroat | 44 | 54 | 6 | 11 | 8 | 20 |
| Yellow Warbler | 49 | 57 | 11 | 23 | 8 | 31 |
| Eastern Towhee | 21 | 25 | 13 | 24 | 5 | 8 |
| Northern Cardinal | 41 | 46 | 13 | 25 | 2 | 13 |
| Indigo Bunting | 30 | 34 | 10 | 17 | 10 | 27 |
| all ten species | 412 | 491 | 99 | 180 | 120 | 369 |

**Figure 4.**
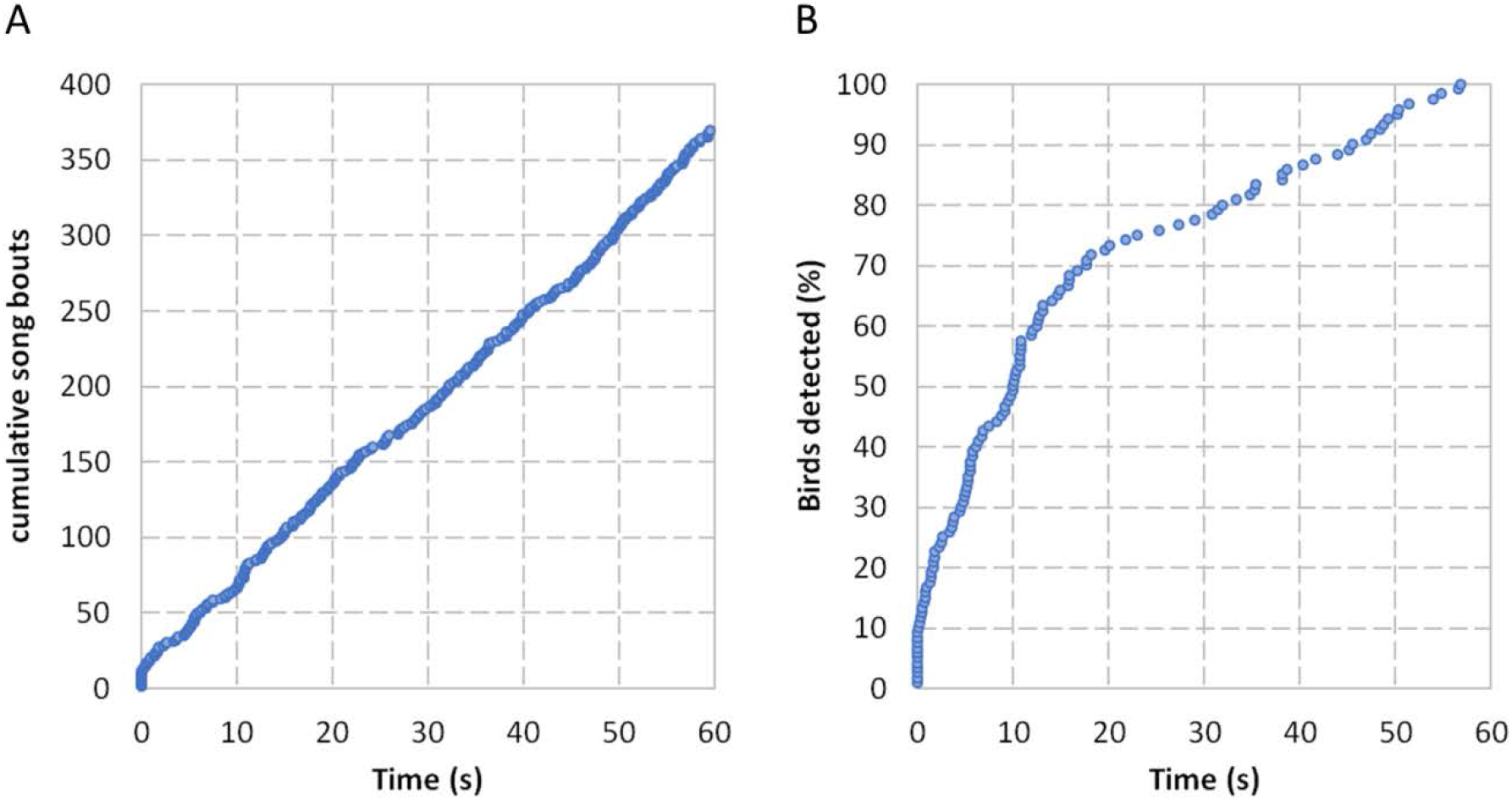
A: Cumulative detections of song bouts from airborne counts, across all 34 counts. B: Cumulative percentage of detections of individual birds from airborne counts across all 34 counts.

The overall patterns of detections from airborne counts were similar to those from terrestrial counts (Figure 5); there was evidence of declining detection rates at distances of greater than 60 m on both point count surveys. Total song detection were higher on airborne counts than in the first minute of terrestrial counts, but lower than the number of individuals detected in 5 minutes (Table 1). Distance sampling derived density estimates for the two most numerous species— Field Sparrow and Song Sparrow—were higher on airborne counts than during the first minute of terrestrial counts, and very comparable with those of five-minute point counts (Figure 6). Density estimates of singing males were lower for both point count methods than the estimated densities of territories derived from mapping (Figure 6). The effective detection radii for Field and Song Sparrows were similar for the airborne counts and terrestrial counts (Table 2).

**Figure 5.**
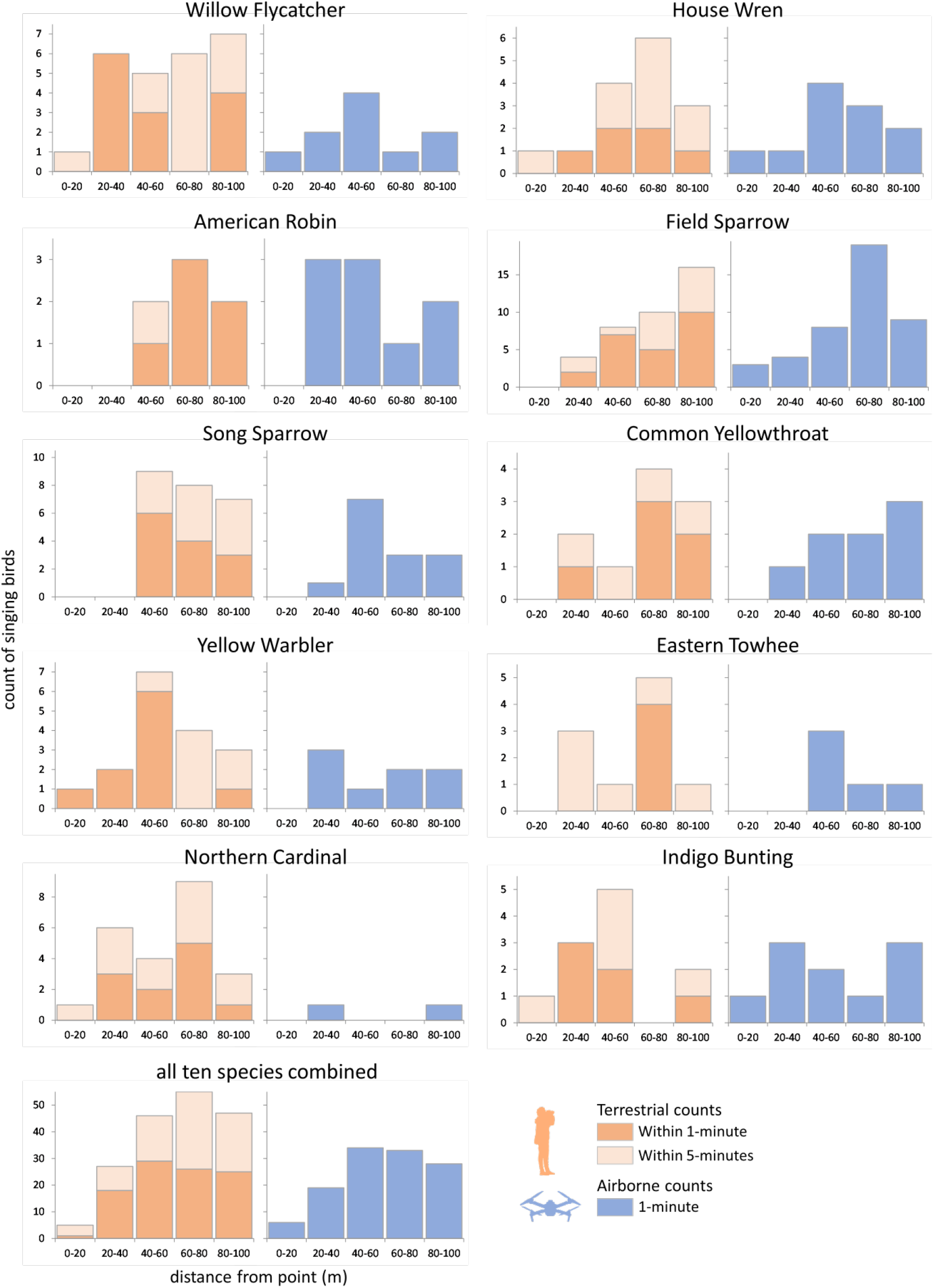
Terrestrial and airborne detections of singing birds of 10 species, across 34 point counts, by distance from point count locations.

**Figure 6.**
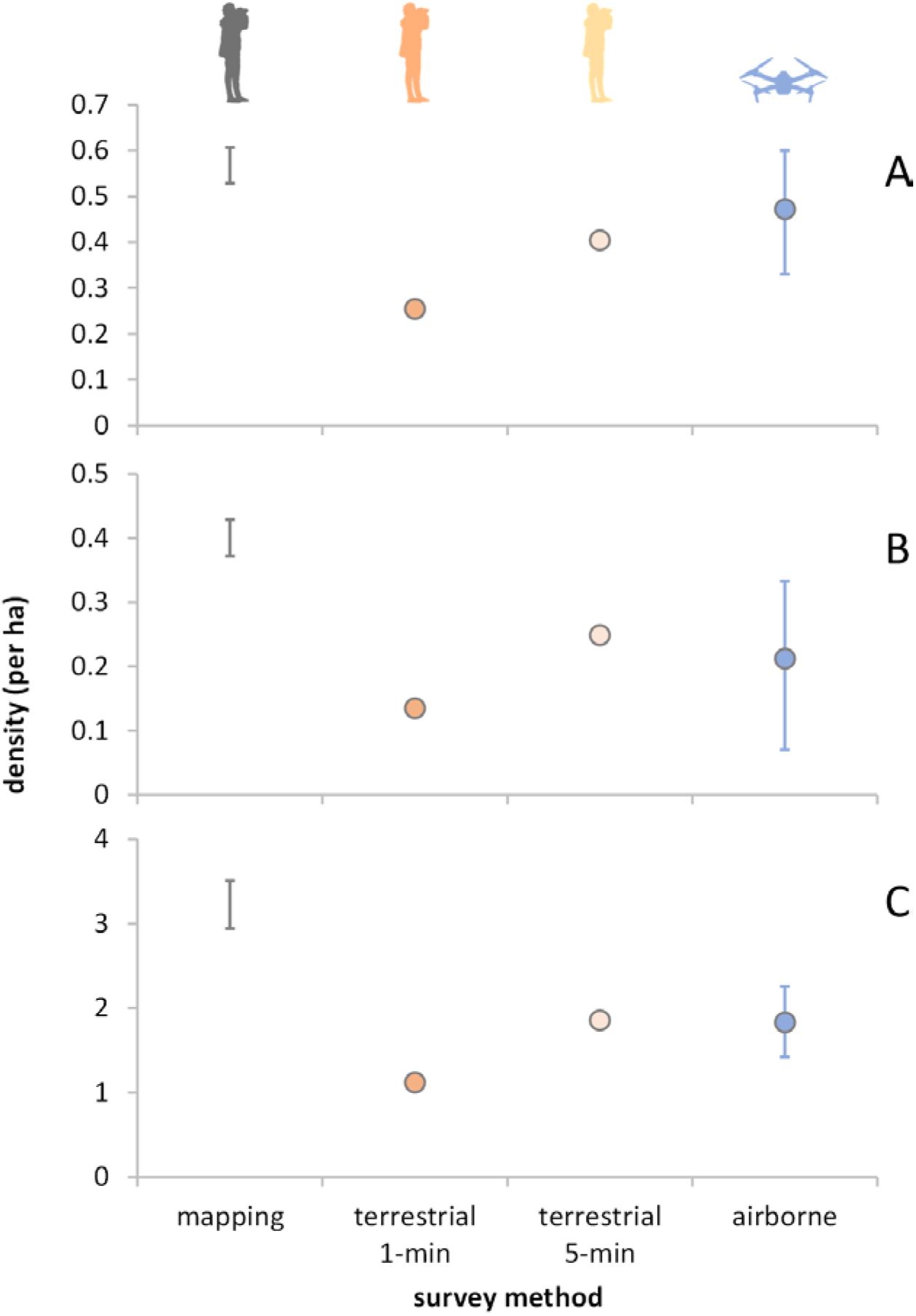
Comparison of density estimates from mapping (territories) and terrestrial and airborne counts (singing birds) for A: Field Sparrow, B: Song Sparrow, and C: all ten species combined. Error bars for mapping shows the low and high estimates of territories. Error bars for estimates from airborne counts are 95% CI. CIs were not calculable for density estimates derived from count estimates.

**Table 2.** Estimated detection radii (meters), calculated using R package *Rdistance*.

|  | terrestrial | airborne |
| --- | --- | --- |
| Field Sparrow | 78.3 | 92.4 |
| Song Sparrow | 90.1 | 90.0 |
| all 10 species combined | 95.0 | 90.0 |

For six of the 10 species the 1-minute airborne counts picked up more singing birds than the first minute of the terrestrial count, and for three species, the airborne counts were higher than after the full 5-minutes of terrestrial counts (Table 1; Figure 5). There was a significant difference in the species composition of detections from airborne counts compared to 5-minute terrestrial counts (χ^2^_9_ = 22.8, *P* = 0.007), with relatively more Field Sparrows (post hoc tests, *P* = 0.045), and fewer Northern Cardinals (post hoc tests, *P* = 0.016) on airborne counts.

## DISCUSSION

Our initial tests prove the concept that it is possible to estimate distances to vocalizing birds with a pair of synchronized recorders suspended from a drone. Our estimates of error are very modest compared to the average observer error of 19 m estimated in a field study by (Alldredge et al. 2007), but we acknowledge that our experimental test was under controlled conditions.

Our field test showed that airborne counts were able to produce data suitable for density estimation that was similar, overall, to that obtained by an experienced fieldworker conducting ground-based point counts. It is important to note that some species appear to have been under-detected on airborne counts, notably the Northern Cardinal. Although our data did not show a decline in song output during the 1-minute drone flights, it is possible that some individual birds stopped singing as the drone approached. Interestingly, in a previous study of seven song bird species, the Northern Cardinal was found to be the most sensitive to drone noise (Wilson et al. 2020). However, as drone technology has matured, small quadcopters have become steadily quieter, and smaller, and further technical developments (e.g. Hioka et al. 2019) could result in drones that are quiet enough to greatly reduce, if not eliminate, noise disturbance effects.

That our density estimates from point counts were very different to those derived from territory mapping was not a surprise; differences, which appear to vary by species, habitat, and specific of protocols, have been widely reported (Shankar Raman 2003, Howell et al. 2004, Newell et al. 2013). The advantages of point counts (either terrestrial or airborne) over mapping is that they don’t require access to an entire study area (Gregory et al. 2004), and they are generally much more time-efficient. Even though our mapping study was scaled back to just four visits to each part of the study area, it required 66 hours of fieldwork, and many hours of collating and analyzing maps.

There are several potential sources of error in our airborne count distance estimates. One of the main sources of error is whether the altitude of the recorders is accurate, and how high off the ground the vocalizing bird is perched (i.e. accuracy of distances *a* and *b*). Drone GPS accuracy is now exceptional, with the manufacturer of the DJI Mavic claiming vertical accuracy of 0.1 m (https://www.dji.com/mavic/info), so we assume that error due to GPS accuracy is trivial. However, the fact that the height of a bird from the ground is not known could introduce significant imprecision and bias into distance estimates. Our method allows estimated perching heights of a particular species from the ground to be incorporated into calculations. Heights could be measured by field validation or estimated using expert opinion. Even so, we currently recommend that our technique is only valid for birds that reliably vocalize from the ground or in low vegetation, and hence is potentially useful for birds of open habitats, including grasslands, wetlands, bare ground, and shrub/scrub. We caution that over-estimating the perching height of birds would lead to an under-estimate of bird densities, because it would be assumed that the bird was further away than it actually was (see Figure 3B).

Another assumption of our technique is that birds vocalize from a horizontal plane. Uneven topography and slopes would therefore introduce inaccuracy into distance estimates, the extent of which merit further investigation. We note that in the case of even and modest slopes, upslope and downslope errors should balance, hence distance estimates will be less precise, but not necessarily biased. Applying our technique in topographically complex locations, where errors could include significant biases, would be more challenging.

A more general limitation of airborne bioacoustics is that drone noise on recordings may mask low frequency songs and calls (Wilson et al. 2017). For such species only those closest to the drone are likely to be detected, reducing the effective sampling distance of this survey method, which could reduce sample sizes. There are, of course, other complications inherent with using drones for survey work including ethics safety, the requirement for additional training and possibly licensing for drone pilots, and the need for relatively calm weather conditions (Linchant et al. 2015, Wallace et al. 2018). However, we have not found any of these to inhibit our ability to use these techniques in Pennsylvania, USA.

As with more general use of ARUs, airborne bioacoustics has some clear disadvantages when compared with traditional bird counts techniques. The most important is that recorders do not pick up visual detections, and hence will undercount species that rarely vocalize when compared with counts by a field ornithologist. In some instances, it may be difficult to be certain how many individual birds of each species are audible in a recording, but airborne count using our method have an advantage over ARUs – the ability to estimate distances provides an additional clue when the analyst is trying to decipher how many birds are detected on a recording.

### Future developments

Importantly, our airborne counts were of just one minute duration, showing that rapid density estimation may be possible using drones. Further, as 74% of singing bird detections were within the first 20 seconds, we believe that it may be possible to use our method to conduct a rapid series of airborne counts in a very short period of time. For example, it would be feasible to fly two adjacent drone missions, each with up to 15 x 30-second counts spaced every 200 m, for a total of 30 point counts densely covering an area of 1.2 km^2^ in point counts in just one hour of field work. This highly efficient data gathering makes it more feasible to repeat point counts over a day or season, which potentially allows for more robust population estimates using occupancy models (Royle and Nichols 2003, Hayes and Monfils 2015) or spatially replicated N-mixture models (Royle 2004). An added advantage of using drones for aerial bioacoustics surveys under those circumstances is that they are highly replicable, both in terms of location and duration (due to the ability to precisely program missions), and because audio recordings provide a permanent record of bird song that can be analyzed by multiple observers after the fact (Shonfield and Bayne 2017).

Although our analysis shows that drones could be a very efficient way to collect data on songbird abundance, it is important to consider the extra analytical time that the method would entail, when compared to terrestrial counts. It took more than one hour of data analysis per airborne count to identify each song bout and then estimate the distance to the bird using the TDoA method – but note that we did localize every song bout, whereas it may be only necessary to localize the first detection.

In addition to trying very rapid assessment with short duration point counts, our technique could easily be modified to fly airborne line transects, which would be even more time-efficient. Two potential pitfalls of that approach are that the recorders may not suspend vertically from a travelling vehicle, and drone noise increase with speed – even a slow-moving quadcopter is noticeably noisier than one hovering.

Recent advances have shown that the distances of vocalizing birds from automated recording units can be estimated from relative sound levels (Yip et al. 2019), which could also be measured using airborne bioacoustics. Combining TDoA based distance estimates with those derived from measurements of sound levels offers an intriguing avenue for technique validation, and for potentially developing a robust technique that combines estimates from the two techniques to reduce uncertainty.

This initial study used off-the-shelf and inexpensive prosumer-level drones and recorders. Results might be greatly improved with custom-designed drones, to reduce drone noise, and custom-built audio recorders. A recorder with the ability to simultaneously record two tracks from input lines of different lengths, would negate the need for manually synchronizing the tracks from the two recorders. We were not able to find a lightweight recording device that satisfied those requirements, but there are options that would be suitable for larger drones that are capable of carrying more payload. However, larger drones are noisier, so the trade-off between payload and the potential for noise disruption needs careful consideration.

## Conclusions

We show that it is feasible to estimate distances of vocalizing birds from a drone, which therefore allows robust estimation of abundance in the same way that traditional bird surveys do. This technique could be applied to places that are difficult or dangerous for point count technicians to access. Further, our study shows that gathering bird abundance data this way could be highly efficient, so this technique may be of broader interest. While the technique outlined here is subject to various assumptions, and will likely not work for all species or habitats, we conclude that with further development of equipment and analytical methods, airborne bioacoustics could provide a useful new way to estimate the abundance of vocal bird species.

## Acknowledgements

We thank Alyssa Kaewwilai and Francesca Garrison for help testing fieldwork protocols. Our study site is on the land of the Susquehannock peoples.

## Funding statement

This work was supported by the Cross-Disciplinary Science Institute at Gettysburg College (X-SIG) and endowed funds from the Cargill Foundation.

## Ethics statement

The drone was flown with the permission of Gettysburg College, under restrictions placed by the Federal Aviation Authority’s Part 107 Remote Pilot License. The use of State Game Lands 249 was with granted permission of the Pennsylvania Game Commission.

## Author Contributions

A.M.W. formulated the research question. M.A.I., P.S.O., L.B.S, M.D.S. and A.M.W. developed fieldwork protocols and conducted the fieldwork. D.B.G. developed analytical methods. All authors contributed to writing and editing the manuscript.

## Notes

### Competing Interest Statement

The authors have declared no competing interest.

## Literature Cited

Afán, I., M. Máñez, and R. Díaz-Delgado (2018). Drone Monitoring of Breeding Waterbird Populations: The Case of the Glossy Ibis. Drones 2:42.

Alldredge, M. W., T. R. Simons, and K. H. Pollock (2007). A Field Evaluation of Distance Measurement Error in Auditory Avian Point Count Surveys. Journal of Wildlife Management 71:2759–2766.

August, T., and T. Moore (2019). Autonomous drones are a viable tool for acoustic bat surveys. bioRxiv:673772.

Betts, M. G., D. Mitchell, D. A. W., and J. Bêty (2007). Uneven Rates of Landscape Change as a Source of Bias in Roadside Wildlife Surveys. Journal of Wildlife Management 71:2266.

Bibby, C. J. (Colin J.., Ecoscope Applied Ecologists., British Trust for Ornithology., Royal Society for the Protection of Birds., and BirdLife International. (2000). Bird census techniques. Academic Press.

Bioacoustics Research Program (2014). Raven Pro: Interactive Sound Analysis Software. The Cornell Lab of Ornithology, Ithaca, New York.

Borrelle, S. B., and A. T. Fletcher (2017). Will drones reduce investigator disturbance to surface-nesting birds? Marine Ornithology 45:89–94.

Buckland, S. T., D. R. Anderson, K. P. Burnham, J. L. Laake, S. T. Buckland, D. R. Anderson, K. P. Burnham, and J. L. Laake (2005). Distance Sampling. In Encyclopedia of Biostatistics. John Wiley & Sons, Ltd, Chichester, UK.

Campbell, M., and C. M. Francis (2013). Using Stereo-Microphones to Evaluate Observer Variation in North American Breeding Bird Survey Point Counts. http://dx.doi.org/10.1525/auk.2011.10005.

Christie, K. S., S. L. Gilbert, C. L. Brown, M. Hatfield, and L. Hanson (2016). Unmanned aircraft systems in wildlife research: Current and future applications of a transformative technology. Frontiers in Ecology and the Environment 14:241–251.

Darras, K., P. Batáry, B. Furnas, A. Celis-Murillo, S. L. Van Wilgenburg, Y. A. Mulyani, and T. Tscharntke (2018). Comparing the sampling performance of sound recorders versus point counts in bird surveys: A meta-analysis. Journal of Applied Ecology 55:2575–2586.

Ebbert, D. (2019). chisq.posthoc.test: a post hoc analysis for Pearson’s chi-squared test for count data. [Online.] Available at https://cran.r-project.org/package=chisq.posthoc.test.

Fu, Y., M. Kinniry, and L. N. Kloepper (2018). The Chirocopter: A UAV for recording sound and video of bats at altitude. Methods in Ecology and Evolution 9:1531–1535.

Gregory, R. D., D. W. Gibbons, and P. F. Donald (2004). Bird census and survey techniques. In Bird Ecology and Conservation. Oxford University Press, pp. 17–56.

Hioka, Y., M. Kingan, G. Schmid, R. McKay, and K. A. Stol (2019). Design of an unmanned aerial vehicle mounted system for quiet audio recording. Applied Acoustics 155:423–427.

Hodgson, J. C., R. Mott, S. M. Baylis, T. T. Pham, S. Wotherspoon, A. D. Kilpatrick, R. Raja Segaran, I. Reid, A. Terauds, and L. P. Koh (2018). Drones count wildlife more accurately and precisely than humans. Methods in Ecology and Evolution. https://doi.org/10.1111/2041-210X.12974

Howell, C. A., P. A. Porneluzi, R. L. Clawson, and J. Faaborg (2004). Breeding Density Affects Point-Count Accuracy in Missouri Forest Birds / (La densidad de aves anidando afectan la exactitud de los conteos de puntos en un bosque de Missouri). Journal of Field Ornithology 75:123–133.

Kloepper, L. N., and M. Kinniry (2018). Recording animal vocalizations from a UAV: Bat echolocation during roost re-entry. Scientific Reports 8:8–13.

Leitão, P. J., F. Moreira, and P. E. Osborne (2011). Effects of geographical data sampling bias on habitat models of species distributions: a case study with steppe birds in southern Portugal. http://dx.doi.org/10.1080/13658816.2010.531020.

Linchant, J., J. Lisein, J. Semeki, P. Lejeune, and C. Vermeulen (2015). Are unmanned aircraft systems (UASs) the future of wildlife monitoring? A review of accomplishments and challenges. Mammal Review 45:239–252.

Macauley Library. (2014). The Cornell Guide to Bird Sounds: Master Set for North America.

Mennill, D. J., J. M. Burt, K. M. Fristrup, and S. L. Vehrencamp (2006). Accuracy of an acoustic location system for monitoring the position of duetting songbirds in tropical forest. The Journal of the Acoustical Society of America 119:2832–2839.

Miller, D. L., E. Rexstad, L. Thomas, J. L. Laake, and L. Marshall (2019). Distance sampling in R. Journal of Statistical Software.

Mulero-Pázmány, M., S. Jenni-Eiermann, N. Strebel, T. Sattler, J. J. Negro, and Z. Tablado (2017). Unmanned aircraft systems as a new source of disturbance for wildlife: A systematic review. PLOS ONE 12:e0178448.

Newell, F. L., J. Sheehan, P. B. Wood, A. D. Rodewald, D. A. Buehler, P. D. Keyser, J. L. Larkin, T. A. Beachy, M. H. Bakermans, T. J. Boves, A. Evans, et al. (2013). Comparison of point counts and territory mapping for detecting effects of forest management on songbirds. Journal of Field Ornithology 84:270–286.

Nowak, M. M., K. Dziób, and P. Bogawski (2019). Unmanned Aerial Vehicles (UAVs) in environmental biology: A review. European Journal of Ecology 4:56–74.

Pöysä, H., J. Kotilainen, V.-M. Väänänen, and M. Kunnasranta (2018). Estimating production in ducks: a comparison between ground surveys and unmanned aircraft surveys. European Journal of Wildlife Research 64:74.

Shankar Raman, T. R. (2003). Assessment of census techniques for interspecific comparisons of tropical rainforest bird densities: a field evaluation in the Western Ghats, India. Ibis 145:9–21.

Shonfield, J., and E. M. Bayne (2017). Autonomous recording units in avian ecological research: current use and future applications. Avian Conservation and Ecology 12.

Valle, R. G., and F. Scarton (2019). Effectiveness, efficiency, and safety of censusing eurasian oystercatchers haematopus ostralegus by unmanned aircraft. Marine Ornithology 47:81–87.

Wallace, P., R. Martin, and I. White (2018). Keeping pace with technology: drones, disturbance and policy deficiency. Journal of Environmental Planning and Management 61:1271–1288.

Wilson, A. M., J. Barr, and M. Zagorski (2017). The feasibility of counting songbirds using unmanned aerial vehicles. The Auk 134:350–362.

Wilson, A. M., K. S.. Boyle, J. L. Gilmore, C. J. Kiefer, and M. F. Walker (2020). Species-Specific Responses of Bird Song Output in the Presence of Drones. Manuscript submitted for publication. https://doi.org/10.1101/2020.07.19.211045

Yip, D. A., E. C. Knight, E. Haave-Audet, S. J. Wilson, C. Charchuk, C. D. Scott, P. Sólymos, and E. M. Bayne (2019). Sound level measurements from audio recordings provide objective distance estimates for distance sampling wildlife populations. Remote Sensing in Ecology and Conservation. https://doi.org/10.1002/rse2.118

